# Deletion of *MEC1* suppresses replicative senescence of the *cdc13-2* mutant in *Saccharomyces cerevisiae*

**DOI:** 10.1101/2023.02.03.527016

**Authors:** Yue Yao, Enikő Fekete-Szücs, Fernando R. Rosas Bringas, Michael Chang

## Abstract

In *Saccharomyces cerevisiae*, telomerase recruitment to telomeres depends on a direct interaction between Cdc13, a protein that binds single-stranded telomeric DNA, and the Est1 subunit of telomerase. The *cdc13-2* allele disrupts telomerase association with telomeres, resulting in progressive telomere shortening and replicative senescence. The Mec1/ATR kinase is both a positive and negative regulator of telomerase activity, and is required for the cell cycle arrest in telomerase-deficient senescent cells. In this study, we find that deletion of *MEC1* suppresses the replicative senescence of *cdc13-2*. This suppression is dependent on telomerase, indicating that Mec1 antagonizes telomerase-mediated telomere extension in *cdc13-2* cells to promote senescence.

## Introduction

Telomeres are nucleoprotein structures that protect the ends of chromosomes by preventing natural chromosome ends from being recognized and processed as DNA double-strand breaks (Jain and Cooper 2010). Telomeric DNA consists of G/C-rich DNA repeats, with the G-rich strand extending to form a 3′ single-stranded overhang. Specialized proteins bind to both the double-stranded telomeric repeats and the 3′ single-stranded overhang to ensure proper telomere function. Due to incomplete DNA replication and nucleolytic degradation, telomeres shorten with each cell division. Cells express the enzyme telomerase to extend telomeres and counteract this ″end replication problem″ (Jain and Cooper 2010). In the budding yeast *Saccharomyces cerevisiae*, telomerase minimally consists of the protein catalytic subunit Est2 and the non-coding RNA subunit TLC1 (Lingner et al. 1997; Singer and Gottschling 1994). Additional proteins, Est1 and Est3, are required for telomerase activity in vivo, and are thought to be involved in telomerase recruitment and/or activation (Wellinger and Zakian 2012). Elimination of any of the Est subunits or TLC1 results in gradual telomere shortening and eventual replicative senescence—the so-called ″ever shorter telomeres″ (*est*) phenotype (Lundblad and Szostak 1989; Lendvay et al. 1996; Singer and Gottschling 1994).

Cdc13 is a single-stranded telomeric DNA binding protein that is essential for telomere end protection and telomerase recruitment (Wellinger and Zakian 2012). These functions are separable. The telomerase recruitment function is mediated by a direct interaction between the recruitment domain (RD) of Cdc13 and Est1 (Pennock et al. 2001). The Cdc13 RD contains two conserved Est1-binding motifs called Cdc13_EBM-N_ and Cdc13_EBM-C_ (Chen et al. 2018). Mutations in Cdc13_EBM-N_ abolish the Cdc13-Est1 interaction in vitro, but in vivo cause only a modest reduction in Est1 telomere association and telomere length (Chen et al. 2018). The Cdc13_EBM-N_-Est1 telomerase recruitment pathway works in parallel with a second pathway involving Sir4, the yKu complex, and TLC1. Sir4 is recruited to telomeres via the double-strand telomeric DNA binding protein Rap1 (Moretti et al. 1994). Sir4 interacts with the Yku80 subunit of the yKu complex (Roy et al. 2004), which binds to the tip of a 48-nt hairpin in TLC1 (Peterson et al. 2001; Stellwagen et al. 2003; Chen et al. 2018). Mutations that disrupt the yKu-TLC1 interaction (e.g. *tlc1Δ48*) cause a modest reduction in telomere length (Peterson et al. 2001). Disrupting both the Cdc13_EBM-N_-Est1 and yKu-TLC1 interactions results in an *est* phenotype (Chen et al. 2018).

The role of the Cdc13_EBM-C_ motif is more enigmatic. The well-studied *cdc13-2* allele has a mutation (E252K) located within this motif (Chen et al. 2018), and causes a dramatic reduction in Est1 telomere association in vivo and an *est* phenotype (Chan et al. 2008; Nugent et al. 1996). However, the mutant Cdc13-2 protein can still interact with Est1 in vitro (Wu and Zakian 2011). Thus, although it is clear that the Cdc13_EBM-C_ motif is required for stable association of telomerase to telomeres, it is unclear how exactly it does so (Chen et al. 2018).

The Tel1 and Mec1 PI3K-like DNA damage checkpoint kinases (ATM and ATR in mammals, respectively) are also important for telomerase activity. Strains harboring a mutation in *TEL1* maintain short, but stable, telomeres (Greenwell et al. 1995). Mec1 has only a modest role in telomere length homeostasis, but *mec1 tel1* double mutants have an *est* phenotype (Ritchie et al. 1999), which can be suppressed by expression of either a Cdc13-Est1 or Cdc13-Est2 fusion protein (Tsukamoto et al. 2001). ATM and ATR are also required for telomere elongation in mouse and human cells (Lee et al. 2015; Tong et al. 2015). In addition to this positive regulatory role, Mec1 also inhibits telomerase at DNA double-strand breaks through two separate mechanisms: (1) via direct phosphorylation of Cdc13 on amino acid S306 (Zhang and Durocher 2010), and (2) via phosphorylation of the helicase and telomerase inhibitor Pif1, which requires the Mec1 downstream kinases Rad53 and Dun1 (Makovets and Blackburn 2009). Mec1 is also required for responding to critically short telomeres and activating a G2/M cell cycle arrest in telomerase-null senescent cells (Enomoto et al. 2002; IJpma and Greider 2003; Abdallah et al. 2009).

We recently discovered that mutation of *PIF1* can suppress the *est* phenotype of the *cdc13-2* mutant (Fekete-Szücs et al. 2022). In this study, we show that deletion of *MEC1* can also suppress the *cdc13-2 est* phenotype. Thus, Mec1 acts to inhibit telomerase activity in *cdc13-2* strains. Mec1-dependent phosphorylation of Cdc13 and Pif1 are either not required or insufficient for this inhibition.

## Materials and Methods

### Yeast strains and plasmids

All yeast strains used in this study are listed in Table 1. Standard yeast genetic and molecular methods were used (Sherman 2002; Amberg et al. 2005). Plasmids pFR95 (pRS415-*cdc13-E252K*), pYY1 (pRS415-*cdc13-S306A*), and pYY2 (pRS415-*cdc13-E252K,S306A*) were created by site-directed mutagenesis of pDD4317 (pRS415-*CDC13*; Strecker et al. 2017) using primers designed by NEBaseChanger and the Q5 Site-Directed Mutagenesis Kit (New England Biolabs, cat. no.: E0554S). The mutations were confirmed by DNA sequencing. Plasmid pEFS4 (pRS415-*cdc13-F237A*) was generated in the same manner from a previous study (Fekete-Szücs et al. 2022).

**Table 1.** Yeast strains used in this study.

| Strain name | Genotype | Source |
| --- | --- | --- |
| YYY2 | *MATa/MATα cdc13-2::natMX/CDC13 mec1ΔTRP1/MEC1 sml1ΔHIS3/sml1ΔHIS3 ADE2/ADE2 can1-100/can1-100 his3-11,15/his3-11,15 leu2-3,112/leu2-3,112 trp1-1/trp1-1 ura3-1/ura3-1 RAD5/RAD5* | This study |
| YYY3 | *MATa/MATα cdc13-2::natMX/CDC13 tel1ΔURA3/TEL1 ade2-1/ADE2 can1-100/can1-100 his3-11/his3-11 leu2-3,112/leu2-3,112 trp1-1/trp1-1 ura3-1/ura3-1 RAD5/RAD5* | This study |
| YYY7 | *MATa/MATα cdc13-2::natMX/CDC13 est1ΔkanMX/EST1 mec1ΔTRP1/MEC1 sml1ΔHIS3/sml1ΔHIS3 ADE2/ADE2 can1-100/can1-100 his3-11,15/his3-11,15 leu2-3,112/leu2-3,112 trp1-1/trp1-1 ura3-1/ura3-1 RAD5/RAD5* | This study |
| EFSY133 | *MATa/MATα cdc13-2::natMX/CDC13 mec1ΔTRP1/MEC1 sml1ΔHIS3/SML1 tlc1Δ48::hphMX/TLC1 ade2-1/ADE2 can1-100/can1-100 his3-11,15/his3-11,15 leu2-3,112/leu2-3,112 trp1-1/trp1-1 ura3-1/ura3-1 RAD5/RAD5* | This study |
| YYY81 | *MATa/MATα cdc13ΔhphMX/CDC13 mec1ΔTRP1/MEC1 sml1Δ/SML1 tlc1Δ48::kanMX/TLC1 ADE2/ADE2 can1-100/can1-100 his3-11,15/his3-11,15 leu2-3,112/leu2-3,112 trp1-1/trp1-1 ura3-1/ura3-1 RAD5/RAD5 pRS415-cdc13-F237A* | This study |
| EFSY79 | *MATa/MATα cdc13-2::natMX/CDC13 pif1-m2::URA3-pif1-m1-4A/PIF1 rad52ΔhphMX/RAD52 ADE2/ADE2 can1-100/can1-100 his3-11,15/his3-11,15 leu2-3,112/leu2-3,112 trp1-1/trp1-1 ura3-1/ura3-1 RAD5/RAD5* | This study |
| YYY6 | *MATa/MATα cdc13-2::natMX/CDC13 mec1ΔTRP1/MEC1 rad53ΔkanMX/RAD53 sml1ΔHIS3/sml1ΔHIS3 ADE2/ADE2 can1-100/can1-100 his3-11,15/his3-11,15 leu2-3,112/leu2-3,112 trp1-1/trp1-1 ura3-1/ura3-1 RAD5/RAD5* | This study |
| YYY9 | *MATa/MATα cdc13ΔkanMX/CDC13 ade2-1/ade2-1 can1-100/can1-100 his3-11,15/his3-11,15 leu2-3,112/leu2-3,112 trp1-1/trp1-1 ura3-1/ura3-1* | This study |
|  | *RAD5/RAD5 pRS415-cdc13-E252K* |  |
| YYY10 | *MATa/MATα cdc13ΔkanMX/CDC13 ade2-1/ade2-1 can1-100/can1-100 his3-11,15/his3-11,15 leu2-3,112/leu2-3,112 trp1-1/trp1-1 ura3-1/ura3-1 RAD5/RAD5 pRS415-cdc13-S306A* | This study |
| YYY11 | *MATa/MATα cdc13ΔkanMX/CDC13 ade2-1/ade2-1 can1-100/can1-100 his3-11,15/his3-11,15 leu2-3,112/leu2-3,112 trp1-1/trp1-1 ura3-1/ura3-1 RAD5/RAD5 pRS415-cdc13-E252K,S306A* | This study |
| YYY12 | *MATa/MATα cdc13ΔkanMX/CDC13 pif1-m2::URA3-pif1-m1/PIF1 ade2-1/ade2-1 can1-100/can1-100 his3-11,15/his3-11,15 leu2-3,112/leu2-3,112 trp1-1/trp1-1 ura3-1/ura3-1 RAD5/RAD5 pRS415-cdc13-E252K* | This study |
| YYY13 | *MATa/MATα cdc13ΔkanMX/CDC13 pif1-m2::URA3-pif1-m1-4A/PIF1 ade2-1/ade2-1 can1-100/can1-100 his3-11,15/his3-11,15 leu2-3,112/leu2-3,112 trp1-1/trp1-1 ura3-1/ura3-1 RAD5/RAD5 pRS415-cdc13-E252K* | This study |
| YYY14 | *MATa/MATα cdc13ΔkanMX/CDC13 pif1-m2::URA3-pif1-m1/PIF1 ade2-1/ade2-1 can1-100/can1-100 his3-11,15/his3-11,15 leu2-3,112/leu2-3,112 trp1-1/trp1-1 ura3-1/ura3-1 RAD5/RAD5 pRS415-cdc13-S306A* | This study |
| YYY15 | *MATa/MATα cdc13ΔkanMX/CDC13 pif1-m2::URA3-pif1-m1-4A/PIF1 ade2-1/ade2-1 can1-100/can1-100 his3-11,15/his3-11,15 leu2-3,112/leu2-3,112 trp1-1/trp1-1 ura3-1/ura3-1 RAD5/RAD5 pRS415-cdc13-S306A* | This study |
| YYY16 | *MATa/MATα cdc13ΔkanMX/CDC13 pif1-m2::URA3-pif1-m1/PIF1 ade2-1/ade2-1 can1-100/can1-100 his3-11,15/his3-11,15 leu2-3,112/leu2-3,112 trp1-1/trp1-1 ura3-1/ura3-1 RAD5/RAD5 pRS415-cdc13-E252K,S306A* | This study |
| YYY17 | *MATa/MATα cdc13ΔkanMX/CDC13 pif1-m2::URA3-pif1-m1-4A/PIF1 ade2-1/ade2-1 can1-100/can1-100 his3-11,15/his3-11,15 leu2-3,112/leu2-3,112 trp1-1/trp1-1 ura3-1/ura3-1 RAD5/RAD5 pRS415-cdc13-E252K,S306A* | This study |

### Liquid culture senescence assay

Liquid culture senescence assays were performed essentially as previously described (van Mourik et al. 2016). Each senescence assay started with diploid strains. Freshly dissected haploid spores were allowed to form colonies on YPD agar plates after two days of growth at 30°C. Cells from these colonies were serially passaged in liquid culture medium at 24-h intervals. For each passage, the cell density of each culture was measured by optical density (calibrated by cell counting using a haemocytometer), and the cultures were diluted back into fresh medium at a cell density of 2 × 10^5^ cells/ml. Cell density was plotted as a function of population doublings.

### Telomere Southern blot

Telomere length analysis by Southern blotting was performed essentially as previously described (van Mourik et al. 2018). Southern blots were probed with a telomere-specific probe (5′-TGTGGGTGTGGTGTGTGGGTGTGGTG-3′).

## Results and Discussion

### Deletion of *MEC1* can bypass the *est* phenotype of the *cdc13-2* mutant

To study the role of Mec1 in *cdc13-2* cells, we sporulated *CDC13/cdc13-2 MEC1/mec1Δ sml1Δ/sml1Δ* diploid cells and followed the growth of the haploid meiotic progeny by serial propagation in liquid cultures for several days. The *sml1Δ* mutation was necessary to overcome the lethality of *mec1Δ* (Zhao et al. 1998). We find that *cdc13-2 mec1Δ sml1Δ* strains do not senesce (Fig 1A) and maintain short, but stable, telomeres (Fig 1B). Since Mec1 and Tel1 have partially redundant functions, we tested whether deletion of *TEL1* could suppress the *cdc13-2 est* phenotype. We find that it cannot (Fig 1C). The suppression of senescence by *mec1Δ* is telomerase dependent, since it has been previously reported that *mec1Δ sml1Δ tlc1Δ* triple mutants exhibit an *est* phenotype (Enomoto et al. 2002; IJpma and Greider 2003). Nevertheless, since the *cdc13-2* mutation reduces the association of Est1 to telomeres, it was possible that deletion of *MEC1* specifically bypasses the need for Est1 for telomerase activity. To test this idea, we sporulated an *EST1/est1Δ MEC1/mec1Δ sml1Δ/sml1Δ* diploid strain and monitored the growth of the haploid meiotic progeny (Fig 1D). We find that *est1Δ mec1Δ sml1Δ* strains senesce, indicating that deletion of *MEC1* cannot bypass the need for Est1.

**Figure 1.**
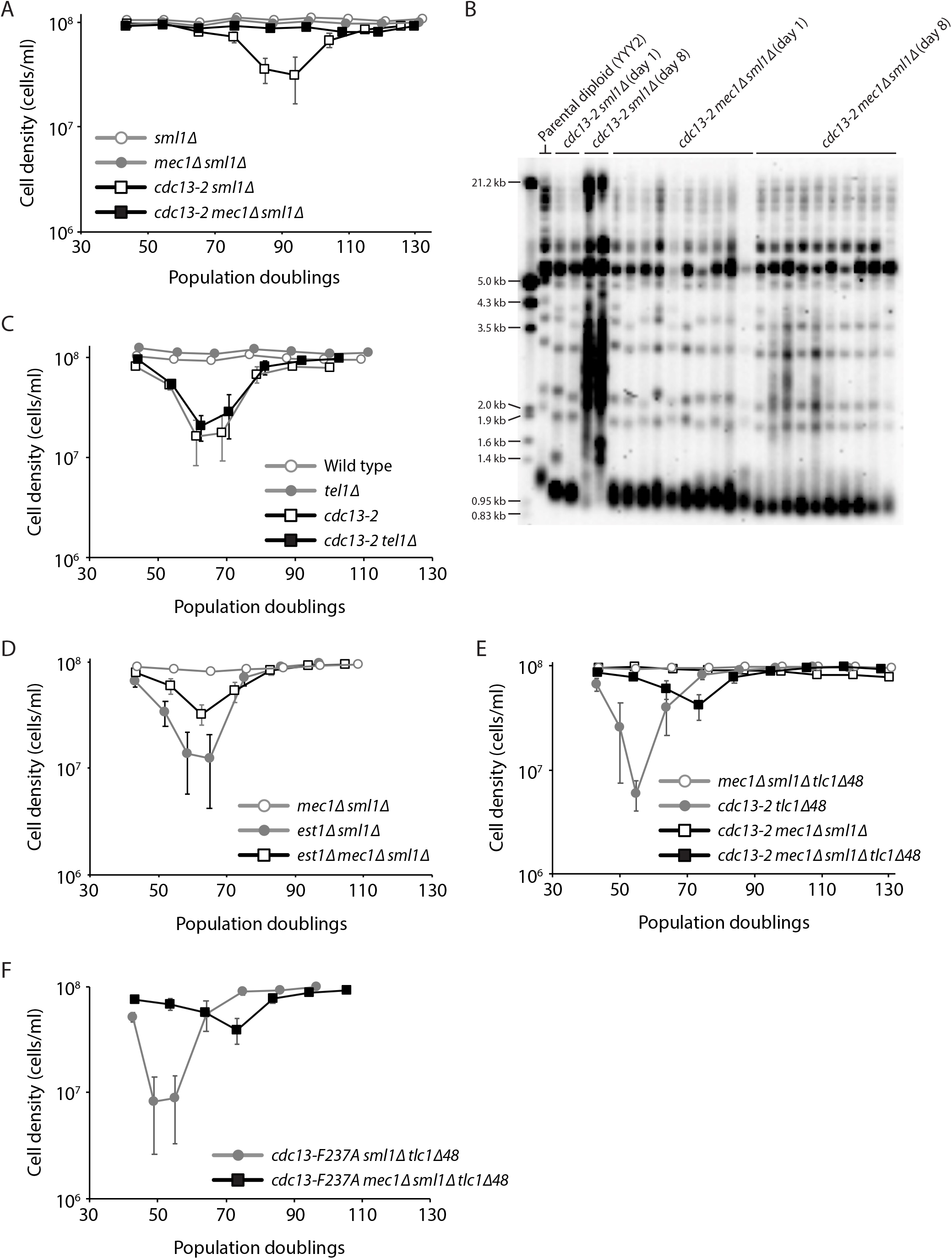
Deletion of *MEC1* suppresses the *cdc13-2 est* phenotype. Senescence was monitored by serial passaging of haploid meiotic progeny derived from the sporulation of YYY2 (A), YYY3 (B), YYY7 (D), EFSY133 (E), and YYY81 (F). Average cell density ±SEM of 3-8 independent isolates per genotype is plotted. (B) Telomere Southern blot analysis of strains of the indicated genotypes.

We previously reported that the *pif1-m2* mutant, which is depleted for nuclear Pif1 (Schulz and Zakian 1994), can suppress the *cdc13-2 est* phenotype, but not if the Sir4-yKu-TLC1 pathway is disrupted (Fekete-Szücs et al. 2022). To test whether this was also true for *mec1Δ* suppression of *cdc13-2*, we monitored the growth of haploid meiotic progeny derived from the sporulation of a *CDC13/cdc13-2 MEC1/mec1Δ SML1/sml1Δ TLC1/tlc1Δ48* diploid strain (Fig 1E). We find that *cdc13-2 mec1Δ sml1Δ tlc1Δ48* strains senesce, suggesting that recruitment of telomerase via the Sir4-yKu-TLC1 pathway is important for telomere maintenance in *cdc13-2 mec1Δ sml1Δ* cells. However, the *tlc1Δ48* mutation leads to a ∼48% reduction in TLC1 abundance (Zappulla et al. 2011), which could also explain the senescence of *cdc13-2 mec1Δ sml1Δ tlc1Δ48* strains. Regardless, our findings indicate that telomere length homeostasis of the *cdc13-2 mec1Δ sml1Δ* strain depends on telomerase. We also observe that *cdc13-2 tlc1Δ48* strains senesce more rapidly than *cdc13-2 mec1Δ sml1Δ tlc1Δ48* strains, which is consistent with Mec1 being important for inducing senescence (Enomoto et al. 2002; IJpma and Greider 2003).

We also previously reported that the *pif1-m2* can suppress the senescence of a *cdc13-F237A tlc1Δ48* double mutant (Fekete-Szücs et al. 2022), in which the Cdc13_EBM-N_-Est1 and yKu-TLC1 interactions are both disrupted (Chen et al. 2018). Unlike *pif1-m2*, we find that *mec1Δ* cannot suppress *cdc13-F237A tlc1Δ48* senescence, although it does delay senescence (Fig 1F).

### Mec1-dependent phosphorylation of Pif1 and Cdc13 is either not require or insufficient to induce senescence in *cdc13-2* cells

Since Pif1 is phosphorylated in a Mec1-dependent manner to inhibit telomerase activity at DNA double-strand breaks (Makovets and Blackburn 2009), and removal of nuclear Pif1 with the *pif1-m2* allele can suppress the *cdc13-2 est* phenotype (Fekete-Szücs et al. 2022), we tested whether the *pif1-4A* mutant, which abrogates Mec1-dependent phosphorylation of Pif1 (Makovets and Blackburn 2009), can also suppress the *cdc13-2 est* phenotype. We monitored the growth of *cdc13-2 pif1-4A* strains derived from the sporulation of a *CDC13/cdc13-2 PIF1/pif1-4A* diploid (Fig 2A). We observe that *cdc13-2 pif1-4A* double mutants senesce. Thus, abolishing Mec1-dependent phosphorylation of Pif1 cannot suppress, or is insufficient for suppressing, the *cdc13-2 est* phenotype.

**Figure 2.**
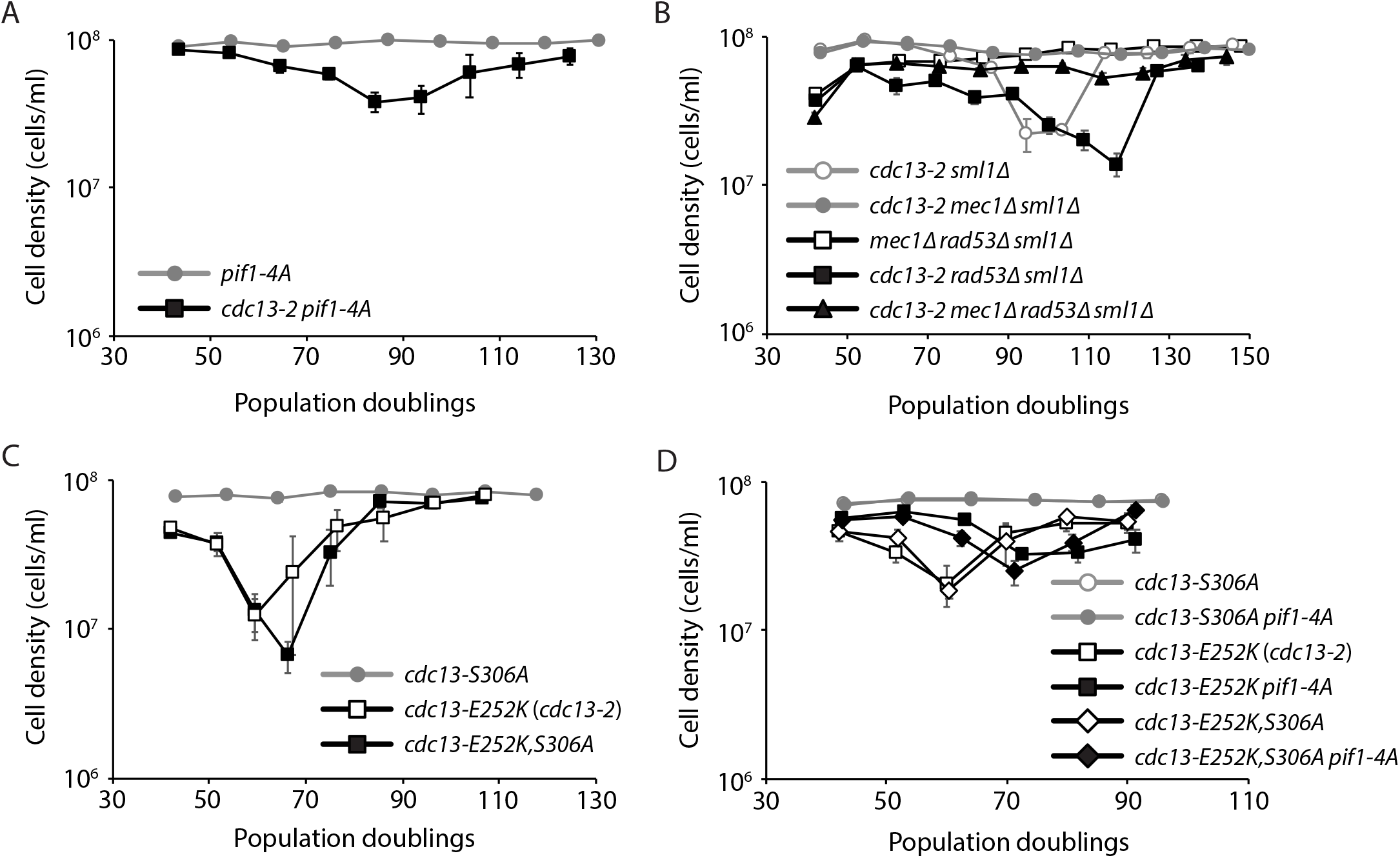
Mec1-dependent phosphorylation of Cdc13 and Pif1 is either not required or insufficient to induce senescence in *cdc13-2* cells. Senescence was monitored by serial passaging of haploid meiotic progeny derived from the sporulation of EFSY79 (A), YYY6 (B), YYY9-11 (C), and YYY12-17 (D).

Mec1 carries out most of its functions through phosphorylation of the effector kinase Rad53 in response to DNA damage or replication stress (Pardo et al. 2017). We therefore tested whether deletion of *RAD53* can suppress the *cdc13-2 est* phenotype. We sporulated a *CDC13/cdc13-2 RAD53/rad53Δ sml1Δ/sml1Δ* diploid strain and monitored the growth of the haploid meiotic progeny (Fig 2B). Deletion of *RAD53* was unable to suppress the *cdc13-2 est* phenotype, indicating that Mec1-dependent activation of Rad53 is either not required or insufficient for inducing senescence in *cdc13-2* cells.

Mec1 can phosphorylate Cdc13 on amino acid S306, in a Rad53-independent manner, and this phosphorylation is important to inhibit telomerase activity at DNA double-strand breaks (Tseng et al. 2006; Zhang and Durocher 2010). To test whether the *cdc13-S306A* mutation could suppress the *cdc13-2 est* phenotype, we combined the *cdc13-2* mutation (E252K) with the S306A mutation. We find that *cdc13-E252K,S306A* mutants senesce (Fig 2C). To rule out the possibility that Mec1-dependent phosphorylation of both Cdc13 and Pif1 are important to induce senescence in *cdc13-2* cells, we monitored the growth of *cdc13-E252K,S306A pif1-4A* cells (Fig 2D). While the *pif1-4A* mutation delays senescence of *cdc13-E252K* and *cdc13-E252K,S306A* cells, it cannot suppress senescence. Thus, Mec1-dependent phosphorylation of Cdc13 and Pif1 are either not required or insufficient for inducing senescence in *cdc13-2* cells. Our findings are consistent with previous observations suggesting that Mec1-dependent phosphorylation of Cdc13 and Pif1 do not inhibit telomerase-mediated extension of native telomeres, only at DNA double-strand breaks (Zhang and Durocher 2010; Makovets and Blackburn 2009).

At present, we do not know what is the relevant Mec1 target or targets responsible inducing senescence in *cdc13-2* cells. Since the *tlc1Δ48* mutation causes *cdc13-2 mec1Δ sml1Δ* cells to senesce, it is possible that Mec1 inhibits the Sir4-yKu-TLC1 telomerase recruitment pathway. Sir4 is among many putative targets of Mec1 identified using mass spectrometry-based approaches (Smolka et al. 2007; Chen et al. 2010; Bastos de Oliveira et al. 2015). Further studies are needed to fully elucidate the role of Mec1 in regulating telomerase activity.

## Acknowledgments

Y.Y. was supported by an Abel Tasman Talent Program scholarship. F.R.R.B. was supported by a CONACYT scholarship.

